# Parallel processing of working memory and temporal information by distinct types of cortical projection neurons

**DOI:** 10.1101/2021.02.16.431447

**Authors:** Jung Won Bae, Huijeong Jeong, Chanmee Bae, Hyeonsu Lee, Se-Bum Paik, Min Whan Jung

## Abstract

It is unclear how different types of cortical projection neurons work together to support diverse cortical functions. We examined the discharge characteristics and inactivation effects of intratelencephalic (IT) and pyramidal tract (PT) neurons—two major types of cortical excitatory neurons that project to cortical and subcortical structures, respectively—in the medial prefrontal cortex of mice performing a delayed response task. We found that IT neurons, but not PT neurons, convey significant working memory-related signals. We also found that the inactivation of IT neurons, but not PT neurons, impairs behavioral performance. In contrast, PT neurons convey far more temporal information than IT neurons during the delay period. Our results indicate a division of labor between IT and PT projection neurons in the prefrontal cortex for the maintenance of working memory and for tracking the passage of time, respectively.

## INTRODUCTION

The cerebral cortex is populated by multiple types of excitatory and inhibitory neurons wired together in complex circuits. Cortical projection neurons can be grouped into intratelencephalic (IT), pyramidal tract (PT), and corticothalamic (CT) projection neurons, with each group differing from one another in developmental origin, morphology, laminar distribution, and input/output connectivity^1-5^. IT neurons are distributed across all layers of the cortex. They project to other cortical areas as well as bilaterally to the striatum. PT neurons are restricted to layer 5, projecting to ipsilateral subcortical structures. CT neurons are located in layer 6 and they project primarily to the thalamus. Clarifying how the different types of cortical projection neurons contribute to the diverse functions of the cortex is essential for understanding the neural circuit mechanisms that underlie those cortical functions.

Few studies have compared the functions of different types of cortical projection neurons in mice. In the primary motor cortex, IT neurons represent the deviation between current and subsequent behaviors, while PT neurons specifically encode the direction and speed of a subsequent behavioral response^6^. Similarly, PT neurons in the anterior lateral motor cortex—the secondary motor cortex^7^—selectively encode and drive contralateral action selection. IT neurons, in contrast, show mixed directional selectivity with little contralateral bias^8^. These findings suggest that, in motor cortical areas, PT neurons are more involved in the control of specific movements than IT neurons. In the primary visual cortex, PT neurons are more direction-selective and prefer faster stimuli, but exhibit broader tuning for orientation and spatial frequency compared to IT neurons^9-11^. In the primary somatosensory cortex, activity of PT neurons, but not IT neurons, is correlated with perception of whisker movement^12^. In the medial prefrontal cortex (mPFC), optogenetic silencing of CT neurons, but not corticocortical IT neurons, impairs choice flexibility^13^. These studies demonstrate functional differentiation among IT, PT, and CT neurons in different cortical regions. Not only are there few published studies on this subject, most of those that have been published targeted the sensory/motor cortices. Thus, our understanding of the ways different types of cortical projection neurons contribute to the great variety of cortical functions, especially high-order cognitive functions, remains limited.

In the present study, we investigated the roles prefrontal cortical IT and PT neurons play in working memory. Working memory refers to the system by which the brain temporarily holds and manipulates information^14^. Although working memory is essential for an array of complex cognitive tasks, the cortical circuit processes that support working memory remain unclear. We are particularly interested in whether and how the different types of cortical projection neurons contribute to the maintenance of working memory. To investigate this, we examined the discharge characteristics and inactivation effects of IT and PT neurons in the mPFC of mice performing a working memory task.

## RESULTS

### mPFC inactivation impairs behavioral performance

We trained head-fixed mice in a delayed match-to-sample task (Fig. 1a,b). The mice were rewarded with water (3 μl) when, after a delay period, they chose the same water lick port that was presented during the sample phase before the delay period. We found well-trained wild-type (WT) mice (n = 6) that received a bilateral infusion of muscimol into the mPFC (Fig. 1c; histological identification of cannula locations in Supplementary Fig. S1) showed a significantly reduced correct response rate (delay duration, 4 s) with no effect on their lick rate as compared to no-infusion and artificial cerebrospinal fluid (ACSF)-infusion controls (one-way repeated measures ANOVA, correct response rate, *F*(2,10) = 36.121, *p* = 2.7×10^−5^; Bonferroni’s post-hoc test, muscimol versus ACSF, *p* = 0.004; muscimol versus no-infusion, *p* = 0.005; ACSF versus no-infusion, *p* = 0.590; lick rate, *F*(2,10) = 0.474, *p* = 0.636; Fig. 1d). This result clearly implicates the mPFC in this match-to-sample task.

**Fig. 1.**
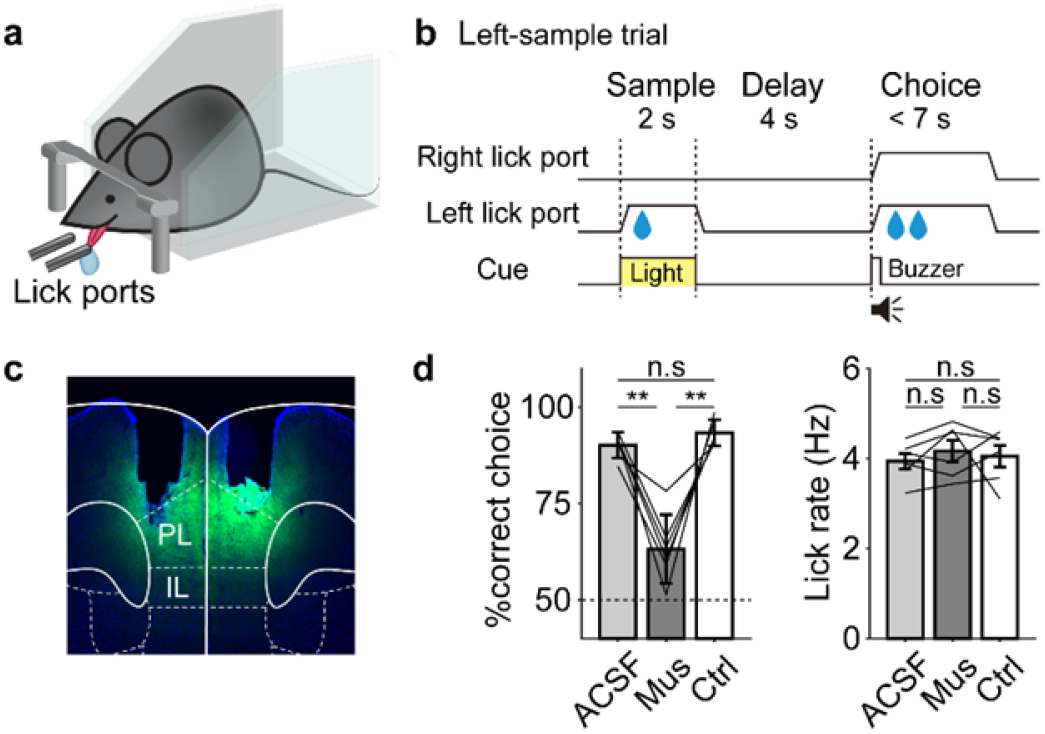
mPFC inactivation impairs behavioral performance. **a**, The experimental setting. Head-fixed mice performed a delayed match-to-sample task. **b**, Schematic for a left-sample trial. Mice obtained a water reward by choosing the lick port presented during the sample phase before a delay period. **c**, A coronal brain section showing cannula tracks and the spread of fluorescein (green) in the mPFC. PL, prelimbic cortex; IL, infralimbic cortex. **d**, Behavioral performance (mean ± SEM across 6 WT mice, left) and lick rate (right) following ACSF, muscimol (Mus), and no-drug (Ctrl) infusions. Thin lines, individual animal data. \*\**p* < 0.01; n.s., non-significant (one-way repeated measures ANOVA followed by Bonferroni’s post-hoc tests).

### Identification and classification of IT and PT neurons

Rxfp3-Cre and Efr3a-Cre knock-in mice were used to target deep-layer IT and PT neurons, respectively^15^ (see Supplementary Fig. S2 for histological verification of IT and PT neurons). We expressed channelrhodopsin-2 (ChR2; Fig. 2a) and implanted in the mPFC an optic fiber for optogenetic tagging and eight movable tetrodes for recording neuronal activity. We recorded 860 and 713 neurons from the mPFC (prelimbic and infralimbic cortices) of 14 Rxfp3-Cre and 12 Efr3a-Cre mice, respectively, during task performance (histological identification of tetrode and optic fiber locations in Supplementary Fig. S1). Then, we classified these neurons as putative pyramidal (PP) and fast-spiking (FS) neurons based on their firing rates and spike waveforms (Fig. 2b). We optically tagged 42 PP, 19 FS, and five unclassified neurons in Rxfp3-Cre mice, as well as 47 PP and 15 FS neurons in Efr3a-Cre mice (Fig. 2b,c; light responses of all optically tagged PP and FS neurons are shown in Supplementary Fig. S3). None of the Cre-expressing neurons expressed parvalbumin (PV; Fig. 2d), a marker for fast-spiking inhibitory interneurons^16^. Nevertheless, to be conservative, we included only optically tagged PP neurons in the main analyses. Their mean discharge rates were 1.9 ± 2.1 in Rxfp3-Cre mice and 1.9 ± 2.5 Hz in Efr3a-Cre mice (mean ± SD).

**Fig. 2.**
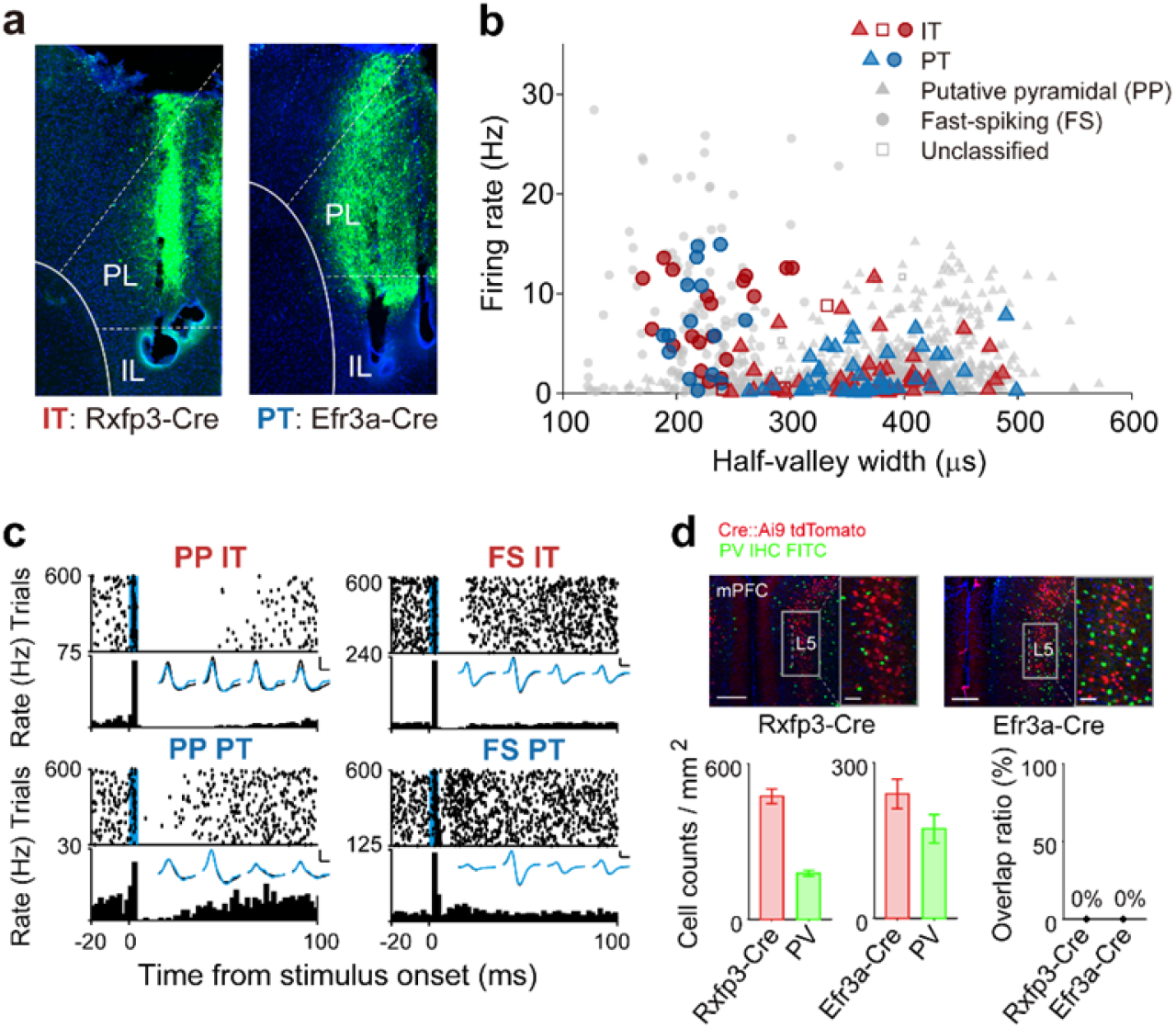
Optical tagging and unit classification. **a**, Sample coronal brain sections showing tetrodes tracks and ChR2-eYFP expression (green). **b**, Neurons were classified as PP (filled triangles) or FS (filled circles) neurons (open squares, unclassified neurons) based on physiological characteristics. Red and blue, optogenetically tagged IT and PT neurons, respectively (gray, untagged neurons). **c**, Sample responses of optically-tagged IT and PT neurons to light stimulation. Top, spike raster plots; each row is one trial and each dot is one spike. Bottom, peri-stimulus time histogram. Time 0 denotes the onset of 5-ms light stimulation (blue bar). The averaged spike waveforms of spontaneous (black) and optically driven (blue) spikes recorded through four tetrode channels are also shown (inset). Calibration, 250 ms and 50 μV. **d**, Top, sample coronal brain sections from Rxfp3-Cre (left) and Efr3a-Cre (right) mice showing tdTomato (red, crossed with Ai9, JAX 007909) and PV (green, immunostained) expression. Note that there is no overlap between Cre-tdTomato and PV expression. Scale bar, left, 200 μm; right, 50 μm. Bottom, group data (mean ± SEM across six brain sections from two animals).

### IT, but not PT, neurons convey working memory-related signals

We subjected the Rxfp3-Cre and Efr3a-Cre mice to two consecutive blocks of the delayed match-to-sample task, one with fixed (4 s; 60-80 trials) and the other with variable (1-7 s; 60-80 trials) delay durations. We reported the discharge characteristics of untagged PP and FS neurons during the fixed and variable delay conditions using a subset of the data^17^. Here, we focus on comparing IT and PT neuronal activity during the fixed delay condition (similar conclusions were obtained from the variable delay condition; Supplementary Fig. S4) using only correct trials, unless otherwise noted.

We found IT, but not PT, neurons tend to show delay-period activity that varies according to sample identity (Fig. 3a). In the group analysis, neural decoding of sample identity using delay-period (4 s) neuronal ensemble activity improved as a function of ensemble size for the IT neurons, but remained similar to the level of chance (50%) across all tested PT neuron ensemble sizes (Fig. 3b). For instance, at ensemble size of 25 neurons (vertical dashed line in Fig. 3b), the neural decoding of sample identity was significantly above the level of chance for IT, but not PT neurons (*t*-test, IT, *t*_99_ = 19.498, *p* = 1.1×10^−35^; PT, *t*_99_ = -0.590, *p* = 0.556; IT vs. PT, *t*_198_ = 14.143, *p* = 7.6×10^−32^; Fig. 3c). This cannot be attributed to the mPFC neurons of Rxfp3-Cre mice carrying stronger working memory-related signals than those of Efr3a-Cre mice (Supplementary Fig. S5). Thus, IT, but not PT, neurons carry significant working memory-related signals during the delay period.

**Fig. 3.**
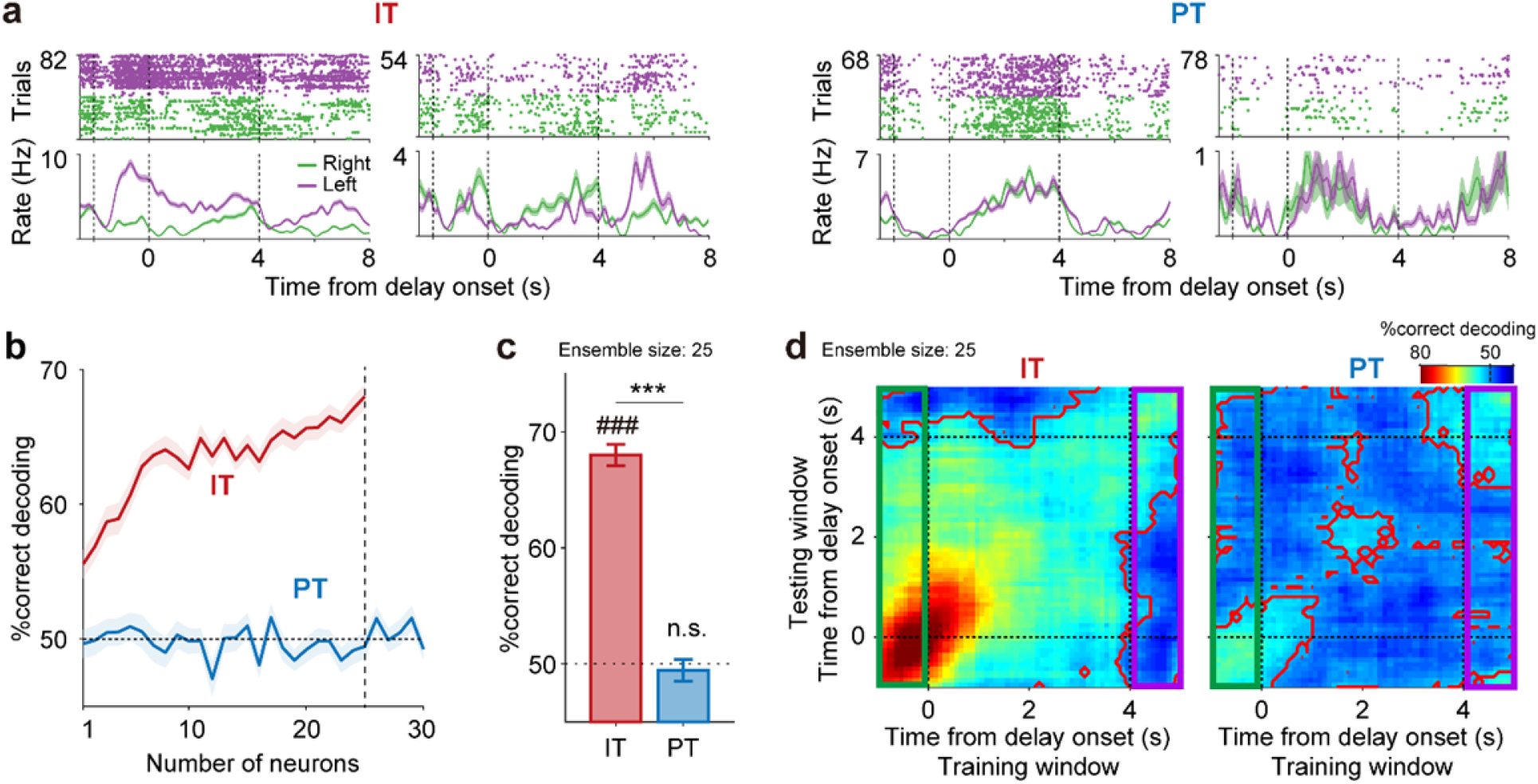
IT, but not PT, neurons convey working memory-related signals. **a**, Sample IT and PT neuronal responses during the task (correct trials only). Top, spike raster plots; bottom, spike density functions (σ = 100 ms). Green, right-sample trials; purple, left-sample trials. **b**, Neural decoding of sample identity as a function of ensemble size. Shading, SEM across 100 decoding iterations. **c**, Decoding performance (mean ± SEM across 100 decoding iterations) at ensemble size of 25 neurons (dashed line in b). ###*p* < 0.001 (above chance level, *t*-test); \*\*\**p* < 0.001 (IT versus PT, *t*-test). **d**, Heat maps for cross-temporal population decoding (n = 25 neurons; 1-s bin advanced in 0.1-s steps). Red contours indicate significant decoding above chance level (*p* < 0.05, *t*-test). Green and purple rectangles denote cross-temporal decoding using sample- or choice-phase neural activity as the training data, respectively.

We then performed a cross-temporal decoding analysis^18,19^ to examine dynamics of sample/choice-dependent neural activity during the delay period (ensemble size = 25 neurons; Fig. 3d). We found that IT neurons show stronger sample-dependent activity than PT neurons during the sample phase (comparison of sample-identity decoding, last 1 s of the sample phase, *t*-test, *t*_198_ = 20.632, *p* = 3.2×10^−51^). Also, as denoted by the red contours in Fig. 3d, IT, but not PT, neurons showed a broad domain of significant (*p* < 0.05, *t*-test) cross-temporal decoding during the delay period, indicating that IT neurons tended to maintain sample/choice-dependent activity persistently during the delay period. When we used sample-phase ensemble activity for the training data (green rectangles in Fig. 3d), IT neurons yielded above-chance decoding of sample identity largely throughout the delay period. In contrast, PT neurons yielded poor sample identity prediction throughout the delay period (mean delay-period decoding performance using sample-phase (last 1 s) ensemble activity as training data, *t*-test, IT, above chance level, *t*_99_ = 11.458, *p* = 7.5×10^−20^; PT, *t*_99_ = -1.101, *p* = 0.274; IT versus PT, *t*_198_ = 8.885, *p* = 3.9×10^−16^). When we used choice-phase ensemble activity as the training data (purple rectangles in Fig. 3d), we found sample identity prediction was poor throughout most of the delay period, except toward the end. Thus, the mean delay-period decoding performance (training data, ensemble activity during the first 1 s of the choice phase) was not significantly above chance level for both IT (*t*-test, *t*_99_ = 0.361, *p* = 0.719) and PT (*t*_99_ = -1.715, *p* = 0.090) neurons (IT versus PT, *t*_198_ = 1.340, *p* = 0.182). These results suggest IT neurons carry stronger retrospective memory (sample identity) than prospective memory (upcoming choice) signals for the majority of the delay period in our task.

### Stronger coupling of IT than PT neurons to theta-frequency activity

The theta-frequency component of the mPFC local field potential (LFP) has been implicated in hippocampal-prefrontal cortical synchrony and spatial working memory^20,21^. Because we found IT, but not PT, neurons convey significant working memory-related signals, we asked whether IT and PT neuronal spikes show different degrees of coupling to the theta-frequency component of the LFP. We found that both IT (n = 40) and PT (n = 45) neurons show enhanced LFP-spike coherence in the theta band (4–8 Hz) during the delay compared to the baseline period (2-s time period before sample onset; *t*-test, IT, *t*_39_ = 4.852, *p* = 2.0×10^−5^; PT, *t*_44_ = 3.605, *p* = 7.9×10^−4^; Fig. 4a). But the coherence of IT neurons was significantly stronger than that of PT neurons during the delay period (*t*-test, *t*_83_ = 2.261, *p* = 0.026; Fig. 4b,c). There were no significant differences in mean discharge rates or in burst firing (*t*-test, *p*-values > 0.103; Fig. 4d,e). These results suggest IT neurons may contribute to the maintenance of working memory, not only via sample-dependent firing, but also by enhancing synchronous firing at theta frequencies.

**Fig. 4.**
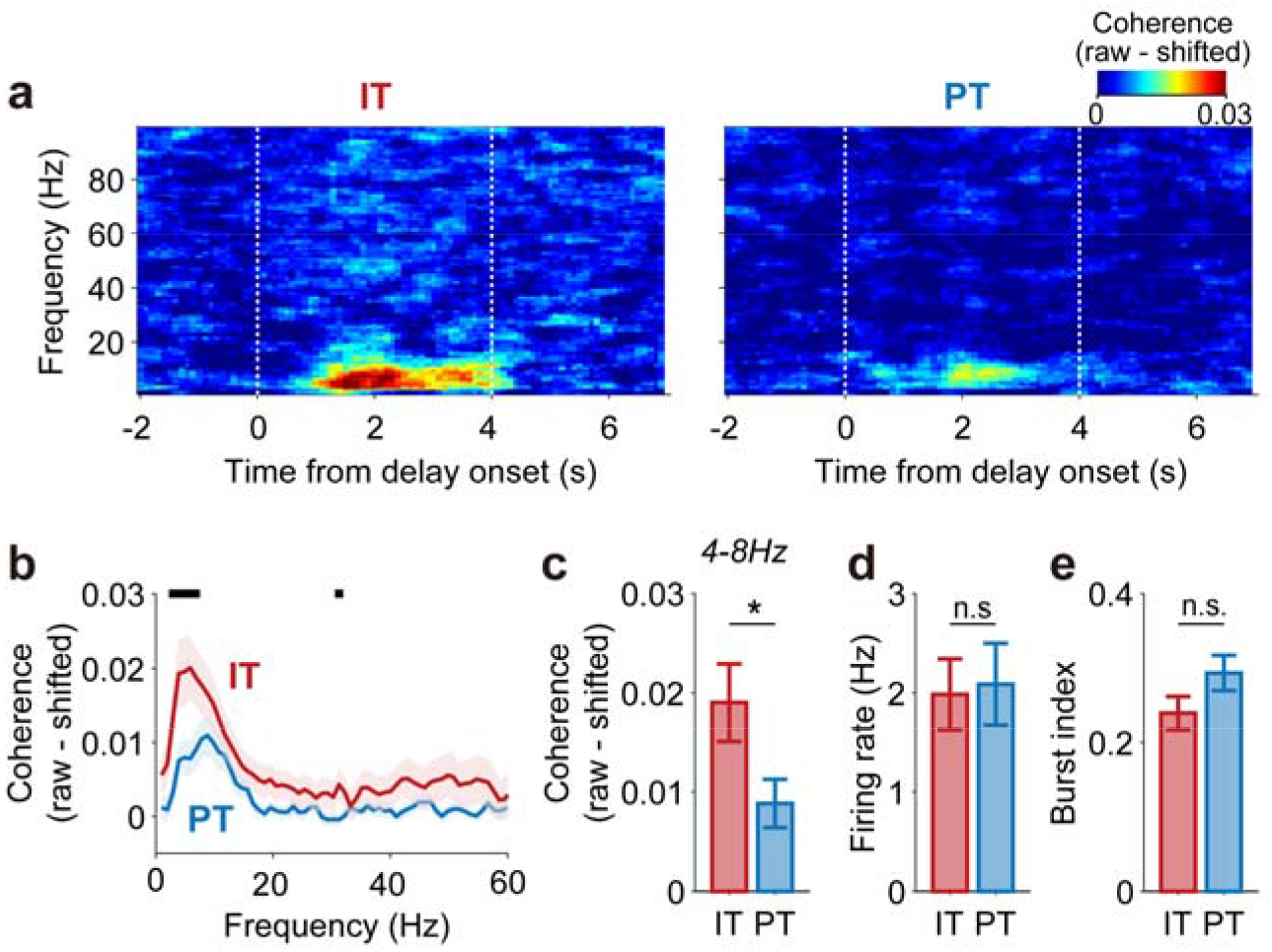
Stronger coupling of IT than PT neuronal spikes to theta-frequency activity. **a**, Frequency-dependent LFP-spike coherence (n = 40 IT and 45 PT neurons). The trial-shifted coherogram was subtracted from the raw coherogram to remove the stimulus-induced coherence. **b**, Mean LFP-spike coherence during the 4-s delay period. Shading, SEM across neurons. Black bars, significant differences between IT and PT (*p* < 0.05, *t*-test). **c-e**, Mean (± SEM across neurons) LFP-spike theta (4-8 Hz) coherences **c**, firing rates **d**, and burst indices (see Methods, **e**) of IT and PT neurons during the 4-s delay period. \**p* < 0.05, *t*-test.

### PT neurons precisely encode the passage of time

Delay-period activity of prefrontal cortex (PFC) neurons conveys not only working memory, but also temporal information^17,22,23^. To assess the temporal information carried by delay-period activity, we divided the delay period into 10 equal bins, leaving out the first 0.5 s to exclude sensory response-related neural activity, leaving a total of 3.5 s. We then decoded bin identity based on IT or PT neuronal ensemble activity to construct a normalized decoding probability map^24^; actual bin number versus predicted bin number; see Methods; Fig. 5a shows an example at ensemble size = 26 neurons). The correlation between actual and predicted bin numbers (*r*) was significantly greater for PT than IT neuronal ensembles (*t*-test, ensemble size = 26 neurons, *t*_198_ = -19.720, *p* = 1.3×10^−48^; Fig. 5b,c). In addition, decoding accuracy (see Methods) was significantly greater for PT than IT neurons (*t*_198_ = -15.594, *p* = 2.7×10^−36^; Fig. 5b,c). These results are not the result of a general tendency of mPFC neurons of Efr3a-Cre mice to carry more temporal information than those of Rxfp3-Cre mice (Supplementary Fig. S6). These findings indicate that PT neurons convey more temporal information than IT neurons during the delay period.

**Fig. 5.**
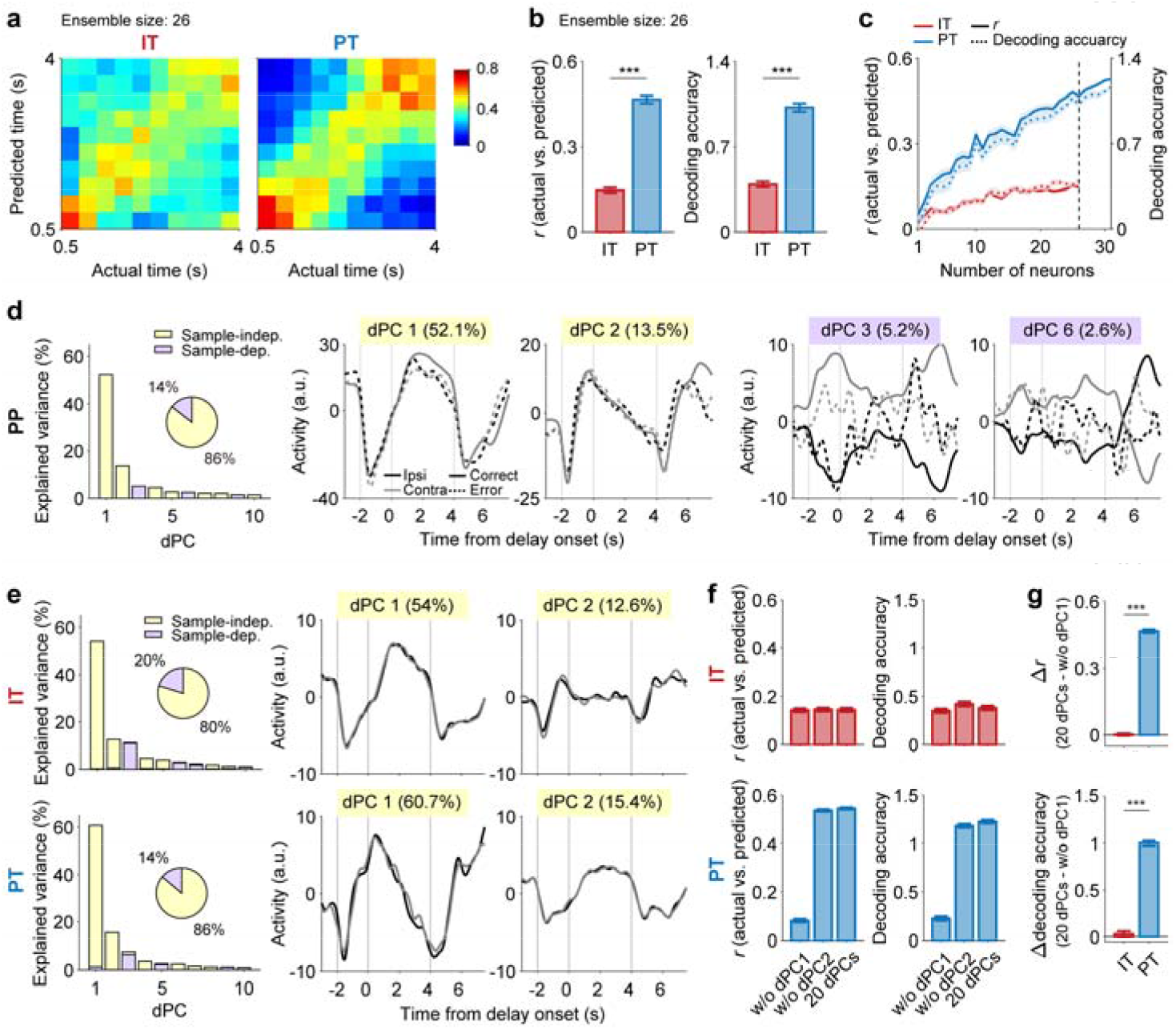
PT neurons precisely encode the passage of time. **a-c**, Neural decoding of time. **a**, The heat maps show normalized decoding probabilities (actual versus predicted bins) averaged across 100 decoding iterations using 26 neurons. **b**, Decoding performances of IT and PT neuronal ensembles (n = 26 neurons; dashed line in c). Left, correlation (*r*) between the actual and predicted bins; right, decoding accuracy (see Methods). \*\*\**p* < 0.001 (*t*-test). **c**, Decoding performance as a function of ensemble size. **d**, Left, variance in neural activity explained by individual dPCs for all analyzed PP neurons (n = 565). Yellow and purple denote sample-independent and sample-dependent dPCs, respectively (see Methods). The pie chart shows the total variance explained by the sample-dependent and sample-independent dPCs. Middle and right, time courses of the top two sample-independent (middle) and sample-dependent (right) dPCs. Solid and dashed lines indicate correct and error trials, respectively. Black and gray denote ipsilateral and contralateral sample trials, respectively. **e**, Left, variance explained by individual dPCs for IT (n = 26) and PT (n = 31) neurons. Right, dPC1 and dPC2. **f**, Temporal decoding using the first 20 dPCs compared with those lacking dPC1 (w/o dPC1) or dPC2 (w/o dPC2). **g**, The amount of decoding performance lost by excluding dPC1 (upper, correlation; lower, decoding accuracy). \*\*\**p* < 0.001 (*t*-test). Results are shown as mean ± SEM across 100 decoding iterations.

To explore characteristics of timing-related mPFC neuronal activity, we performed a demixed principal component analysis (dPCA; analysis time window, [-3.5 8] s from delay onset) using all PP neurons recorded in a session containing at least one error trial for each target and with mean discharge rates ≥ 0.5 Hz (n = 565). We confirmed that the sample-independent demixed principal components (dPCs; those conveying no working memory-related signals) explained the majority of variance, as was previously reported^22,23^. These dPCs showed similar responses between correct and error trials, suggesting that they do not contribute to the animal’s choice of target. This was in contrast to the sample-dependent dPCs that showed different responses between correct and error trials (Fig. 5d; c.f., delay-period components of sample-dependent PT neuronal dPCs did not convey significant working memory-related signals; Supplementary Fig. S7). In the dPCA using PT neurons (n = 31), the first principal component (dPC1) showed ramping activity during the delay period and accounted for ∼60% of neural activity variance. In contrast, there was no monotonic delay-period ramping activity among the first three dPCs of IT neurons (n = 26), although these explained ∼80% of the total variance in neural activity (Fig. 5e). In addition, when dPC1 was removed from the analysis, PT neuronal decoding of the elapse of delay was greatly reduced (Fig. 5f,g). This indicates that PT neuronal temporal information during the delay period was largely conveyed by the ramping activity (Fig. 3a). Collectively, these results indicate that the delay-period activity of IT neurons conveys working memory-related signals with limited temporal information. In contrast, the delay-period activity of PT neurons conveys precise information about the elapsed delay period in the form of ramping activity with minimal working memory-related signal.

### Inactivation of IT neurons impairs working memory

We then examined the effects of selective inactivation of IT or PT neurons on behavioral performance using separate groups of Rxfp3-Cre (n = 12) and Efr3a-Cre (n = 11) mice. We expressed soma-targeted *Guillardia theta* anion-conducting channelrhodopsin-2 (stGtACR2)^25^ and implanted optical probes bilaterally in the mPFC. After training the animals to the performance criterion, we applied continuous laser stimulation during the entire delay period in a randomly selected 30% of 100-200 total trials (Fig. 6a; histological identification of optic fiber locations in Supplementary Fig. S1). We found optical stimulation significantly impaired behavioral performance for trials with 15- and 25-s delays in Rxfp3-Cre mice (two-way repeated measures ANOVA, main effect of laser stimulation, *F*(1,11) = 18.760, *p* = 0.001; main effect of delay duration, *F*(2,22) = 46.117, *p* = 1.3×10^−8^; stimulation×duration interaction, *F*(2,22) = 4.254, *p* = 0.027; Bonferroni’s post-hoc test for stimulation effect, 4-s duration, *p* = 0.727; 15-s, *p* = 0.037; 25-s, *p* = 0.002; Fig. 6b, left), but not in Efr3a-Cre mice (main effect of laser stimulation, *F*(1,10) = 0.003, *p* = 0.959; main effect of delay duration, *F*(2,20) = 6.558, *p* = 0.006; stimulation×duration interaction, *F*(2,20) = 0.094, *p* = 0.911; Fig. 6b, right). Importantly, this occurred without any effect on lick rate or lick response latency (two-way repeated measures ANOVA, no significant main or interaction effects, *p*-values > 0.05; Fig. 6c,d). These results indicate that selective inactivation of IT neurons, but not PT neurons, impairs working memory.

**Fig. 6.**
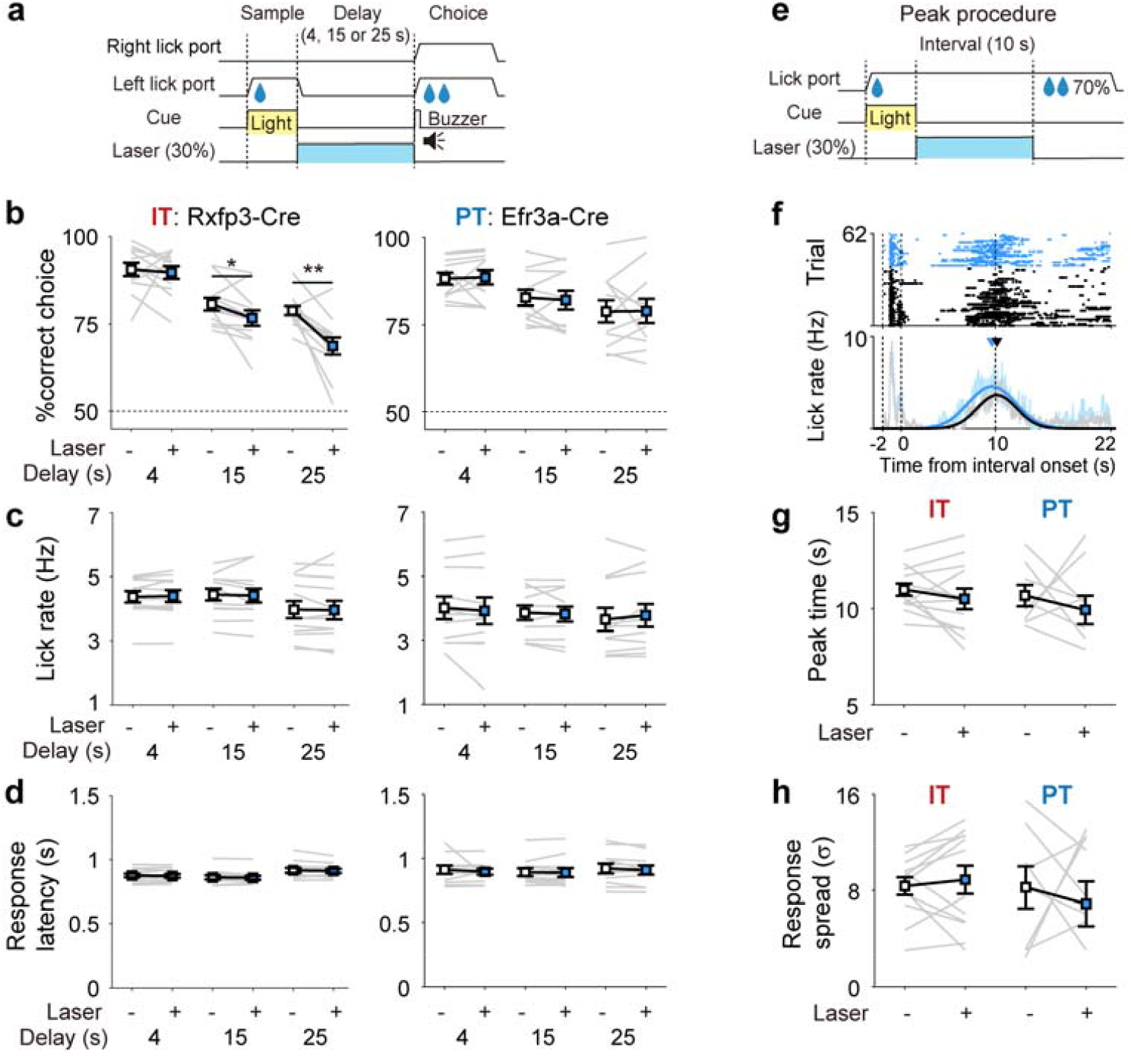
Inactivation of IT neurons impairs working memory. **a**, Schematic for optogenetic inactivation. Laser was given throughout the delay period (blue bar; 4, 15 or 25 s) in a randomly chosen 30% of trials. **b**, Behavioral performance in the absence (-) or presence (+) of laser stimulation for different delay durations (4, 15, and 25 s). \**p* < 0.05; \*\**p* < 0.01 (two-way repeated measures ANOVA followed by Bonferroni’s post-hoc tests). **c**, Lick rate. **d**, Response latency (the latency to first lick since the delay offset). **e**, Schematic for optogenetic inactivation in the peak procedure. **f**, A sample peak procedure session. Only omission trials are shown. Lick raster plots (upper) and lick histograms (bottom) with (blue) or without (gray) laser stimulation are shown. Peak time and SD of the lick response were determined by fitting a Gaussian curve to each lick histogram (thick curves). **g, h** Group data. Peak time and SD of Gaussian-fitted lick responses during omission trials with (+) or without (-) laser stimulation. Gray lines, performances of individual mice; Squares and error bars, mean and SEM.

The lack of any significant effect of IT/PT neuronal inactivation on response latency in the present task may arise from the fact that the delay offset was explicitly signaled by the sound of a buzzer and by the protrusion of the lick ports. We therefore tested the same animals in a peak procedure^26,27^. One lick port delivering water (1.5 μl) was advanced to the animal and a light stimulus (2 s) was presented. Then, after a 10-s interval, water (3 μl) was again delivered in a randomly chosen 70% of trials (Fig. 6e). We also delivered optical stimulation throughout the 10-s interval period in a randomly chosen 30% of trials (Fig. 6e). In this experiment, we did not observe any significant change in the time course of the animal’s licking response (peak time and SD of licking response) upon inactivation of either IT or PT neurons during the reward-omission trials (paired *t*-test, *p*-values > 0.244; Fig. 6f-h). This indicates that neither IT nor PT neurons are indispensable for controlling the timing of the licking response in the present peak procedure.

### Responses of optically-tagged FS neurons differ from those of optically-tagged PP neurons

We subjected only optically-tagged PP neurons to the main analyses because they showed physiological characteristics of typical cortical pyramidal neurons^28,29^ and they out-numbered optically-tagged FS neurons (IT, 42 PP versus 19 FS; PT, 47 PP versus 15 FS). However, that none of the optically-tagged neurons expressed PV raises a possibility that optically-tagged FS neurons might be projection neurons as well. We therefore examined working memory- and timing-related activities of FS IT and FS PT neurons. On the one hand, we found that FS IT, but not FS PT, neurons convey significant working memory-related signals (*t*-test, ensemble size = 14 neuron; FS IT, above chance level, *t*_99_ = 18.471, *p* = 7.6×10^−34^; FS PT, *t*_99_ = -0.574, *p* = 0.567; FS IT versus FS PT, *t*_198_ = 11.859, *p* = 7.4×10^−25^; Fig. 7a,b), which is consistent with the results obtained from the analysis of optically tagged PP neurons. On the other hand, we found that both FS IT and FS PT neurons convey substantial amounts of temporal information and that FS IT neurons convey significantly more temporal information than FS PT neurons (*t*-test, ensemble size = 14 neurons; *r, t*_198_ = 8.150, *p* = 4.1×10^−14^; decoding accuracy, *t*_198_ = 4.059, *p* = 7.1×10^−5^; Fig. 7c-e), which is inconsistent with the results obtained from the analysis of optically tagged PP neurons. The amounts of temporal information conveyed by optically-tagged FS neurons were comparable to that conveyed by PP PT neurons at equivalent ensemble sizes (Fig. 7f,g). In fact, FS IT neurons conveyed significantly larger amounts of temporal information than PP PT neurons, albeit the difference was small (*t*-test, ensemble size = 19 neurons; *r, t*_198_ = 2.273, *p* = 0.024; decoding accuracy, *t*_198_ = 1.703, *p* = 0.090). Thus, optically tagged FS and PP neurons were similar to one another regarding working memory-related neural activity, but different from one another regarding timing-related neural activity.

**Fig. 7.**
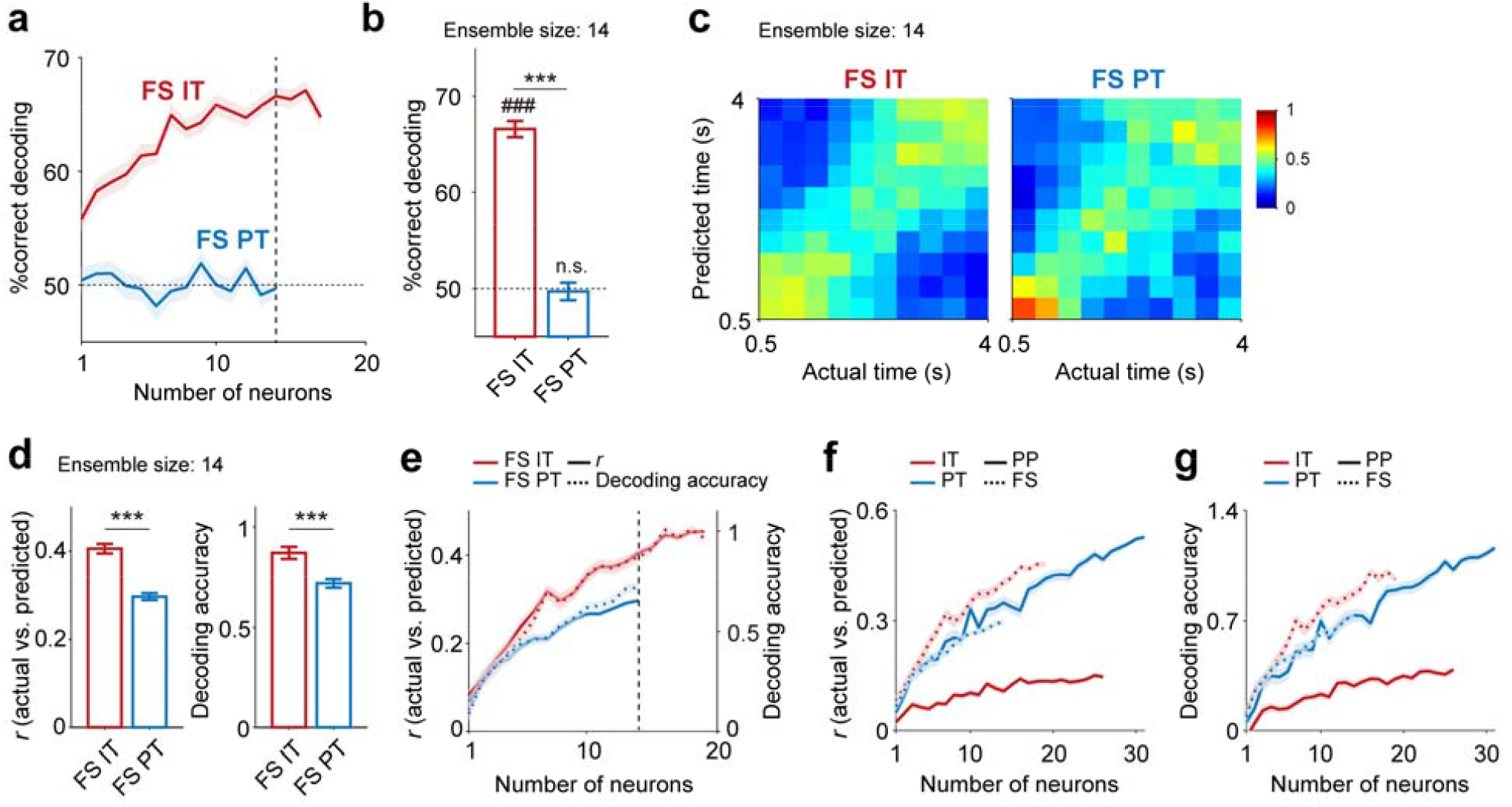
Working memory- and timing-related activity of optically-tagged FS neurons. We repeated the same analyses using optically tagged FS neurons. a, b, Neural decoding of sample identity based on delay-period (4 s) neuronal ensemble activity. **a**, Decoding performance as a function of ensemble size. **b**, Decoding performance at ensemble size of 14 neurons (vertical dashed line in A). ###*p* < 0.001 (above chance level, *t*-test); \*\*\**p* < 0.001 (IT versus PT, *t*-test). The same format as in Fig 3b, c. **c-e**, Timing-related neural activity. **c**, Heat maps showing normalized decoding probabilities (actual versus predicted bins; ensemble size = 14 neurons). **d**, Decoding performance (left, correlation between the actual and predicted bins; right, decoding accuracy) of FS IT and FS PT neuronal ensembles (n = 14, vertical dashed line in E). \*\*\**p* < 0.001 (*t*-test). **e**, Decoding performance as a function of ensemble size. Same format as in Fig. 5a-c. **f, g**, Comparison of temporal decoding across different types of neurons. **f**, Correlation between the actual and predicted bins. **g**, Decoding accuracy. Results are shown as mean ± SEM across 100 decoding iterations.

## DISCUSSION

### Division of labor between IT and PT neurons

Working memory and interval timing are critical components of cognition. Here, we found that IT neurons, but not PT neurons, convey significant working memory-related signals. In contrast, PT neurons carry far more temporal information than IT neurons during the delay period. These results show that, rather than being supported by a common neural substrate^22,30,31^, working memory and interval timing are mediated primarily by two distinct types of projection neurons in the mouse mPFC. This finding may reflect a more generalized functional segregation between neural networks that include IT and PT neurons. For example, the prefrontal cortex may contribute to high-order cognitive functions by sharing working memory-related signals with other cortical areas (IT neuronal network) and to the temporal organization of behavior by sending temporal information to its downstream subcortical structures (PT neuronal network). Given the widespread nature of the cortical neural activity related to working memory and interval timing^32-37^, future studies will be needed to determine whether parallel processing of working memory and temporal information by IT and PT neurons is a general characteristic across the many areas of the cortex.

### Role of IT neurons in the maintenance of working memory

While PT neurons make profuse connections with one another and receive strong projections from IT neurons, IT neurons have sparser interconnections and do not receive projections from PT neurons^2,38-41^. Because profuse interconnections may promote reverberatory activity, PT neurons have been proposed to play a more important role in the maintenance of working memory than IT neurons^40-42^. But inconsistent with this hypothesis, we found IT neurons, rather than PT neurons, convey significant working memory-related signals during the delay period. We also found that the inactivation of IT neurons, rather than PT neurons, impairs behavioral performance. Moreover, IT neurons show stronger coupling to theta frequency oscillations than PT neurons during the delay period. Delay-period theta oscillations and spike-LFP theta synchrony are enhanced in the mPFC in correct trials during working memory tasks^20,43,44^. This enhancement is accompanied by increased hippocampal-mPFC theta synchrony^20,45^ with the level of synchrony showing a positive correlation with behavioral performance^46^. In addition, monosynaptic hippocampal projections to the mPFC preferentially target IT neurons^47^. Together, these results indicate the importance of IT neurons, rather than PT neurons, in working memory.

Previous studies have shown that somatostatin (SST)-expressing interneurons in the mPFC play an important role in the maintenance of working memory. SST neurons, but not PV neurons, show significant sample-dependent activity during the delay period, and the inactivation of SST neurons, but not PV neurons, impairs performance in a spatial working memory task^48^. Also, during probabilistic classical conditioning tasks, SST neurons convey strong expected outcome information until the trial outcome^49^. In addition, SST neurons facilitate hippocampal-mPFC theta synchrony and their inactivation disrupts the directionality of synchrony during a spatial working memory task^21^. These results suggest IT and SST neurons work together to maintain working memory-related signals during the delay period and to regulate the hippocampus-mPFC theta synchrony that facilitates the transfer of spatial information from the hippocampus to the mPFC. It will be important in future studies to investigate how IT and SST neurons work together to support working memory.

### Role of PT neurons in interval timing

The prefrontal cortex is critical for interval timing^50^. We found mPFC PT neurons carry far more temporal information than IT neurons during the delay period. This suggests PT neurons play a major role in keeping track of the passage of time in the mPFC. We have shown previously that the mPFC conveys temporal information largely based on ramping activity^51^. Consistent with this, we found that most PT neuronal temporal information is supported by ramping activity. PT neurons show spike doublets and weaker spike adaptation than IT neurons^39^. In addition, PT neuron pairs show higher paired-pulse ratios and stronger interconnectivity than IT neuron pairs^40^. These facilitatory properties of PT neurons may shape PT neuron ramping activity, but this remains to be tested.

Although PT neurons convey precise temporal information, optogenetic inactivation of PT neurons does not significantly change animal response latency. This indicates PT neurons are dispensable for controlling the timing of the licking response in the present behavioral tasks (delayed match-to-sample and peak procedure tasks). We found that optically-tagged FS (both IT and PT) neurons convey high levels of temporal information during the delay period of the delayed match-to-sample task. It may be that temporal information conveyed by either PT (PP and/or FS) or IT (FS) neurons is sufficient for controlling the timing of the licking response, such that inactivation of one neuron type (IT or PT) is insufficient to alter timing-related behavior in our behavioral tasks. It is also possible that the mPFC plays a more important role in controlling perceptual-timing^51,52^ than motor-timing behavior. These remain unclear.

### Fast-spiking IT and PT neurons

We unexpectedly identified FS IT and PT neurons that do not express PV. The results we obtained from optically-tagged FS and PP neurons were similar to one another with regard to the neural decoding of sample identity (i.e., working memory). They were different from one another, however, with regard to the neural decoding of time; both FS IT and FS PT neurons convey temporal information comparable to that conveyed by PP PT neurons. In fact, FS IT neurons convey slightly, but significantly more temporal information than FS PT and PP PT neurons. This suggests optically-tagged FS and PP neurons represent functionally distinct populations. In this regard, we previously showed that the FS neuronal population conveys quantitative value signals in the mPFC, while optically-tagged PV neurons convey only valence signals, not value signals^49^. This indicates the existence of FS neurons that are functionally distinct from PV neurons in the mPFC. These may correspond to the optically-tagged FS neurons in the present study. It is possible that these optically-tagged FS neurons in the mPFC represent long-range GABAergic neurons^53,54^. But other studies have also identified PV-positive long-range neurons in the mPFC^55,56^. Thus, the exact identity of PV-negative FS neurons must still be clarified. Our results show that FS IT neurons convey both working-memory and temporal information during the delay period. We still need to confirm whether these represent a specific subset of projection neurons that transmit combined working-memory and temporal information. If they do, we also need to identify the downstream targets of their projections.

### IT and PT neurons and neuropsychiatric disorders

The PFC is implicated in diverse neuropsychiatric disorders, such as schizophrenia, attention deficit hyperactivity disorder, obsessive-compulsive disorder, and depression^33,57,58^, all of which are associated with impaired working memory and/or interval timing^59-62^. This raises the possibility that these neuropsychiatric disorders are associated with different patterns of abnormalities in IT versus PT neurons. Thus, a better understanding of how IT and PT neuronal abnormalities in the PFC are associated with the symptoms of various neuropsychiatric disorders should provide important insights into the etiologies of these diseases.

## METHODS

### Subjects

Rxfp3-Cre (MMRRC 036667, Mutant Mouse Resource and Research Center, CA, USA; n = 29) and Efr3a-Cre knock-in (MMRRC 036660; n = 26) mice were used to target deep-layer IT and PT neurons, respectively, in the mPFC. Some mice (14 Rxfp3-Cre and 12 Efr3a-Cre) were used for neurophysiological recordings, some (12 Rxfp3-Cre and 11 Efr3a-Cre) were used for optogenetic inactivation, and the rest (3 Rxfp3-Cre and 3 Efr3a-Cre) were used for verification of IT/PT neurons (Supplementary Fig. S2) or immunohistochemistry (Fig. 2d). An additional 6 WT mice (C57BL/6 J, JAX000664, Jackson Laboratory) were used for muscimol-infusion experiments. The animals were individually housed under a 12 hr light/dark cycle. After 2–3 days of handling, the animals were water-restricted and allowed to drink water only during the tasks. All animal care and experimental procedures were performed in accordance with protocols approved by the directives of the Animal Care and Use Committee of Korea Advanced Institute of Science and Technology (approval number KA2018-08).

### Working memory task

The details of the working memory task are described in our previous study^17^. The animals were trained in a delayed match-to-sample task under head fixation (Fig. 1). Each trial comprised sample, delay, and choice phases. Two water lick ports, located on the left and right sides of the animal, were in the retracted position before trial onset. Each trial began when a randomly chosen lick port (sample) was advanced to the animal and used to deliver a drop of water (1.5 μl). A green LED light located above the sample lick port was illuminated at trial onset. After 2 s, the sample lick port was retracted and the LED light was turned off (onset of delay period). The duration of the delay period was either fixed (4, 15, or 25 s) or variable (1–7 s; uniform random distribution). Both water lick ports were advanced to the animal with a brief (100 ms) buzzer sound at the end of the delay period (onset of the choice phase). If the animal licked the lick port that was presented during the sample period, two drops of water (1.5 μl each) were delivered at the chosen water lick port. Then, both water lick ports were maintained at their advanced positions for 7 s before being retracted (correct trial). Both lick ports were retracted immediately without water delivery if the animal licked the opposite port (incorrect trial). If the animal did not lick either port within 4 s of the delay offset, both lick ports were retracted without water delivery (miss trial; mean ± SEM; 1.8 ± 0.3% in neurophysiological recording sessions and 3.0 ± 0.4% in optogenetic inhibition sessions). A variable inter-trial interval (0–10 s) was then imposed before the next trial began.

All animals were initially trained with a fixed delay (4 s) until they reached the performance criterion (> 70% correct choices for three consecutive daily sessions; 5–10 sessions; 120–160 trials per session). WT mice were then tested under the fixed-delay (4 s) condition with or without drug infusion into the mPFC (in the order of ACSF infusion, muscimol infusion, and no drug infusion for 3 daily sessions). The second group of mice (14 Rxfp3-Cre and 12 Efr3a-Cre) were subjected to unit recordings under the fixed-delay (4 s) condition for 2–4 sessions after reaching the performance criterion. Then, during unit recordings, the mice were trained to perform two consecutive blocks of fixed-delay (4 s; 60–80 trials per block) and variable-delay (1–7 s; 60–80 trials per block) trials (their order reversed across successive sessions). Unit signals recorded during the variable-delay blocks before reaching the performance criterion (1–3 training sessions) were excluded from the analysis. After reaching the performance criterion with the 4-s fixed delay, the third group of mice (12 Rxfp3-Cre and 11 Efr3a-Cre) was subjected to optogenetic inactivation (2 sessions, 100–200 trials per session). This procedure (i.e., training until reaching the performance criterion and then investigating the effects of optogenetic inactivation for 2 sessions) was repeated while increasing the delay duration to 15 s and then to 25 s.

### Peak procedure

Each trial began as the right water lick port was advanced toward the animal, with the left lick port remaining retracted throughout the test. Then, a green LED light located above the advanced lick port was illuminated and a drop of water (1.5 μl) was delivered to animal. The LED light was turned off after 2 s to signal interval onset. The water lick port remained in the advanced position and the animal was allowed to lick freely during the interval phase (10 s). Two drops of water (1.5 μl each) were delivered without any external cue in a randomly chosen 70% of trials at the interval offset. Then, the water lick port was retracted 12 s after the interval offset. The animals were trained for 2–3 days until a sharp increase in anticipatory licking was observed around the interval offset. Three animals that showed low levels of anticipatory licking were excluded from the inactivation experiment.

### Virus injection

A small burr hole (diameter, 0.5 mm) was drilled into the skull (1.84 mm anterior and 0.4–0.5 mm lateral to bregma) under isoflurane (1.5–2.0% [v/v] in 100% oxygen) anesthesia, and a bolus of virus (0.5 μl) was injected 1.25–1.75 mm below the brain surface targeting the mPFC at a rate of 0.05 μl/min. For optogenetic tagging of IT and PT neurons, we injected a double-floxed (DIO) Cre-dependent adeno-associated virus (AAV) vector carrying the gene for channelrhodopsin-2 (ChR2) in-frame and fused to enhanced yellow fluorescent protein (AAV2/2-EF1a-DIO-hChR2(H134R)-eYFP, UNC Vector Core, NC, USA) unilaterally in the left or right mPFC (counterbalanced across animals). For optogenetic inactivation of IT or PT neurons, we injected Cre-dependent AAV carrying soma-targeted *Guillardia theta* anion-conducting channelrhodopsin 2 (pAAV1-hSyn1-SIO-stGtACR2-FusionRed, http://www.addgene.com, plasmid 105677) bilaterally.

### Neurophysiology and optogenetics

A hyperdrive, containing an optic fiber (core diameter, 200 μm) and 8 tetrodes, was implanted unilaterally above the prelimbic cortex (1.84 mm anterior and 0.45 mm lateral to bregma; 1.25 mm ventral to brain surface) immediately after virus injection. The optic fiber was left in the same location, but individual tetrodes were advanced gradually each day (0.05–0.1 mm per day) once unit recording began to record different unit signals across days. For optical tagging of IT or PT neurons, 473 nm laser pulses (5-ms duration; 1 Hz; 0.2, 0.5, and 1 mW; 200 pulses at each intensity; Omicron-Laserage, Rodgau-Dudenhofen, Germany; controlled by Labview, National Instruments, TX, USA) were delivered at the end of each recording session. The majority of neurons were recorded from the prelimbic cortex (numbers of prelimbic versus infralimbic cortical neurons, Rxfp3-Cre mice, 39 versus 3 optically-tagged PP, 18 versus 1 optically-tagged FS, 596 versus 98 untagged PP, and 55 versus 22 untagged FS neurons; Efr3a-Cre, 46 versus 1 optically-tagged PP, 11 versus 4 optically-tagged FS, 497 versus 94 untagged PP, and 30 versus 23 untagged FS neurons) as judged by the locations of tetrode marking lesions and tetrode advancement histories. Unit signals and LFPs were amplified (10,000× and 1000×, respectively), band-pass filtered (600–6000 Hz and 0.1–9000 Hz, respectively), and digitized (32 kHz and 1 kHz, respectively) using the Cheetah data acquisition system (Neuralynx, MT, USA).

For optical inactivation of IT and PT neurons, two optical fibers were implanted above the prelimbic cortex. A continuous 473 nm laser stimulation (9 mW) was delivered in a randomly chosen subset of trials (30% of a total of 100–200 trials) throughout the delay period (4, 15, or 25 s) in the delayed match-to-sample task and throughout the interval phase (10 s) in the peak procedure.

### Muscimol infusion

Two cannulas (diameter, 1 mm) were implanted bilaterally targeting the prelimbic cortex (1.84 mm anterior and 0.4 mm lateral to the bregma; 1.2 mm ventral to brain surface) under isoflurane (1.5–2.0% [v/v] in 100% oxygen) anesthesia. After recovery from surgery (3–5 d) and training in the task with a fixed delay (4 s; 5–10 d), the animal’s performance was tested with ACSF, muscimol, or no drug infusion. ACSF (Tocris Bioscience, Bristol, UK, 0.2 μl) or muscimol (5 mM; Sigma-Aldrich, MO, USA, 0.2 μl) was injected into the prelimbic cortex at a rate of 0.02 μl/min 45 minutes before behavioral testing. Fluorescein (5 mM; Sigma-Aldrich, MO, USA, 0.2 μl) was injected before sacrifice to visualize drug spread in the brain. Histological examination showed the spread of fluorescein mostly in the prelimbic cortex, with a small amount spreading to the upper infralimbic cortex (Fig. 1c).

### Histology

For the mice used for unit recordings, small marking lesions were made by passing electrical current (20 mA, 15 s) through one channel of each tetrode at the end of the final recording session. Coronal sections (40 μm) of the brains were prepared according to a standard histological procedure^63^ and the locations of the marking lesions, electrode tracks, cannula tracks, and optic fiber tracks were determined by examining images obtained (10x) with a Zeiss Axio Scan.Z1 slide scanner (Zeiss, Jena, Germany). In order to examine potential overlaps between PV neurons and optically tagged FS neurons, Rxfp3- and Efr3a-Cre mice were crossed with the Ai9 tdTomato reporter line (JAX 007909, Jackson Laboratory) and PV neurons were immunostained according to the following procedure. Brain slices were incubated with blocking solution (5% goat serum in PBS with 0.2% Triton-X 100) for 1 h at room temperature and then with primary antibodies diluted in the blocking solution overnight (anti-PV primary antibody, ab11427, Abcam, Cambridge, UK; 1:1000) at 4°C. The slices were rinsed with the blocking solution and incubated with secondary antibodies for 2 h at room temperature (FITC-conjugated anti-rabbit IgG, AP132F, Millipore, Darmstadt, Germany; 1:200). The slices were then washed three times with PBS and mounted with 4’,6-diamidino-2-phenylindole (DAPI)-containing Vectashield (Vector Laboratories, CA, USA). Cell counting was done manually on brain sections containing the mPFC. Three brain sections from two animals (total six sections) were quantified for each Cre line.

### Isolation and classification of units

Putative single units were isolated by manually clustering various spike-waveform parameters using the MClust Software (http://redishlab.neuroscience.umn.edu/MClust/MClust.html). Only units with an L-ratio < 0.1 (0.03 ± 0.03; mean ± SD; n = 1573), an isolation distance > 19 (47.1 ± 32.3; mean ± SD), and an inter-spike interval < 2 ms were included in the analysis^64^.

A Gaussian mixture model incorporating mean firing rate, peak-valley ratio, and half-valley width as model parameters was used to classify units into PP and FS neurons^48^. The mean (± SD) firing rate, peak-valley ratio, and half-valley width were 7.3 ± 7.4 Hz, 1.1 ± 0.2, and 216.9 ± 39.0 μs, respectively, for FS neurons (n = 164). For the PP neurons (n = 1374), the same parameters were 1.6 ± 2.3 Hz, 2.3 ± 0.5, and 372.7 ± 57.4 μs, respectively. Only putative PP neurons were included in the analysis unless otherwise noted.

### Identification of IT and PT neurons

Those units that met the following two criteria were considered to be optically tagged neurons. First, the latency of the first spike during the 5-ms window after laser stimulation onset should be significantly (*p* < 0.01) shorter than that in the absence of laser stimulation (a 5-ms window chosen randomly from the 900-ms time period before laser stimulation onset) as determined by both the log-rank test and the stimulus-associated spike latency test^65^. Second, the correlation between laser-driven and spontaneous waveforms should be ≥ 0.8.

### Decoding analysis

Working memory-related neural activity was assessed by decoding the identity of the sample target (left versus right lick port) based on delay-period neuronal ensemble activity using the support vector machine (SVM; ‘fitcsvm’ function of MATLAB, Mathworks Inc., Natick, MA, USA). We combined neural data recorded from different sessions excluding those neurons with mean firing rates < 0.5 Hz or with < 20 correct-choice trials for both left and right targets (n = 17 of 42 IT, 17 of 47 PT, and 25 of 760 untagged PP neurons). We randomly selected a given number of neurons for a given ensemble size and, for each selected neuron, randomly selected 20 correct left-sample and 20 correct right-sample trials from the corresponding fixed-delay (4 s) block. Then a single trial was removed, and sample identity in that trial was decoded based on neuronal ensemble activity in that trial (test trial) and neuronal ensemble activity in the remaining trials (training trials) separated by the sample identity (leave-one-out cross validation). This procedure was repeated 100 times and the percentage of correct decoding was calculated. Per iteration, we used a randomly selected set of IT or PT neurons and, for each neuron, randomly selected sets of 20 left-sample and 20 right-sample trials (out of 25-77 correct left-sample and 26-75 correct right-sample trials). For neural decoding of sample identity in the variable-delay block, we used only those trials with delay durations ≥ 4 s and randomly selected 10 rather than 20 correct trials for each sample target because of a smaller sample size. For cross-temporal decoding^18,19^, we trained the classifier with ensemble activity in a specific time bin (1-s bin advanced in 0.1-s steps) and tested for all time bins spanning 1 s before and after the delay period.

Timing-related delay-period neural activity was assessed by decoding elapsed time based on neuronal ensemble activity using the SVM. We excluded the first 0.5 s of the delay period from this analysis to exclude sensory response-related neural activity. Again, we combined neural data recorded from different sessions except neurons with mean firing rates < 0.5 Hz, randomly selected a set of neurons for a given ensemble size, and randomly selected 20 correct trials for each neuron from the corresponding fixed-delay (4 s) block. We divided the delay period (0.5–4 s; total 3.5 s) into 10 equal duration bins, trained the SVM to predict the bin number (rather than sample identity), decoded the bin number using a leave-one-out cross validation procedure, and peak-normalized the outcome (decoded bin counts) for each training bin to generate a normalized decoding probability map^24^. The decoding performance was estimated in two different ways. First, we calculated Pearson’s correlation between the actual and predicted bin numbers. Second, we calculated the decoding accuracy—the difference in decoding error (the absolute difference between actual and predicted bin numbers) between temporal bin-shuffled data (i.e., the order of 10 temporal bins were randomly shuffled) and the original data. Decoding accuracy measures the degree to which temporal prediction is improved (reduced decoding error) by using the original instead of the temporally shuffled neural data. This procedure was repeated 100 times (using a randomly selected set of neurons and a randomly selected set of 20 correct trials for each neuron per iteration). For temporal decoding in the variable-delay block, we used only those trials with delay durations ≥ 4 s and 10 randomly selected correct trials.

### Spike-LFP coherence

The spike-LFP coherogram, which represents coherence between neural spikes and LFPs, was calculated using the function “cohgramc” in the Chronux toolbox of MATLAB as:

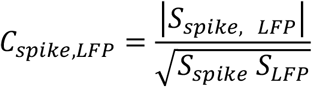

Here, *S*_*spike, LFP*_ is the dot product between the frequency spectrums of spikes and LFP. *S*_*spike*_ and *S*_*LFP*_ were also calculated similarly from the spectrums of spikes and LFPs, respectively. Only the data from fixed-delay trials were included in this analysis. Those neurons whose associated LFP signals reached the ceiling > 10 % of the whole recording duration were excluded from this analysis.

For each neuron, a coherogram was calculated using both the original and the trial-shifted spike trains^66^ in such a way that the trial-shifted coherogram was subtracted from the original coherogram to remove spurious covariation caused by stimulus-locked responses. This procedure was repeated for LFP signals recorded from all tetrodes other than the one detecting a given IT/PT neuron, and the resulting coherograms were averaged across tetrodes. Then, the mean coherogram was averaged across neurons. Mean firing rate and a burst index (the proportion of inter-spike intervals < 100 ms) were also compared between IT and PT neurons to examine any potential effects of firing frequency and firing pattern on coherogram.

### Demixed principal component analysis (dPCA)

We performed a dPCA^23^ to dissociate neural activities related to working memory and temporal information using those neurons recorded in a session containing at least one error trial for each target and with mean discharge rates ≥ 0.5 Hz. A spike density function (σ = 200 ms) spanning the time window 3.5 before and 8 s after delay onset was generated for each trial type (left- and right-sample trials) for each neuron using correct trials. Then, a three-dimensional matrix of neuronal population activity (neuron × time × trial type) was constructed. The matrix was used to demix sample-dependent and sample-independent variances and to obtain PCs as previously described^23^. The first five dPCs explained 86.1, 88.4, and 78.2% of the total variance for IT, PT, and PP population activities, respectively. Spike density functions of left- and right-sample error trials were projected onto the dPC decoder axis when comparing correct versus error trial neural activities (Fig. 5d). To assess the contributions from individual sample-independent dPCs to the representation of temporal information during the delay period, we compared neural decoding of the elapse of delay (see above) by the first 20 dPCs with and without a specific given dPC.

### Statistical tests

Results are presented as mean ± SEM unless noted otherwise. All statistical analyses were performed with MATLAB (version 2017a). One-way repeated measures ANOVA (muscimol effect) and two-way repeated measures ANOVA (laser stimulation effect; group factors: delay duration and laser stimulation) were performed along with Bonferroni’s post-hoc tests to compare the correct response rate, lick rate and response latency in the delayed match-to-sample task. All other comparisons were performed with Student’s *t*-tests. All tests were two-tailed and a *p*-value < 0.05 was used as the criterion for a significant statistical difference unless otherwise specified.

## Data availability

The datasets that support the findings of this study are available from the corresponding author M.W.J. upon request.

## Code availability

MATLAB code used to generate results for this study is available from the corresponding author M.W.J.

## Acknowledgements

This work was supported by the Research Center Program of the Institute for Basic Science (IBS-R002-A1; M.W.J.).

## Author contributions

J.W.B. and M.W.J. designed the study. J.W.B. and H.J. collected and analyzed experimental data. C.B. performed histology. H.L. and S.-B.P. analyzed spike-LFP coherence. J.W.B., H.J., and M.W.J. wrote the manuscript with inputs from all authors. M.W.J. supervised all aspects of the study.

## Materials & Correspondence

Correspondence and requests for materials should be addressed to M.W.J.

