## Supplementary file for "Parallel processing of working memory and temporal information by distinct types of cortical projection neurons"

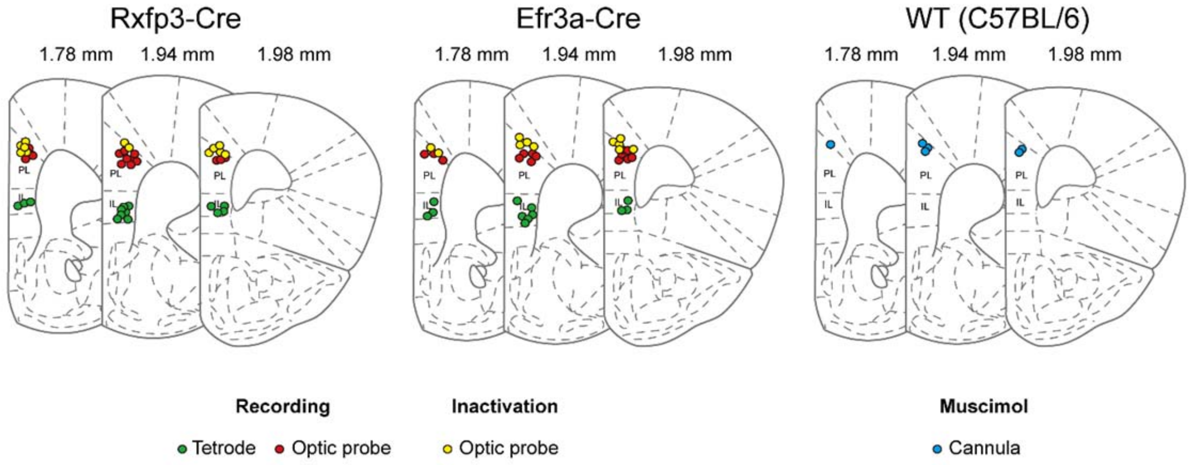


**Supplementary Fig. S1 | Histological identification of cannula, optic probe, and tetrode locations.** Shown are coronal section views of the mouse brain (left to right, 1.78, 1.94, and 1.98 mm anterior to bregma). Circles indicate locations of cannula tips used for muscimol infusion (blue; WT), optical probe tips used for optical tagging (red; Rxfp3-Cre and Efr3a-Cre), tetrode tips at the end of the final recording session for each mouse (green, one representative tetrode for each mouse; Rxfp3-Cre and Efr3a-Cre), and optical probe tips used for inactivation of IT or PT neurons (yellow; Rxfp3-Cre and Efr3a-Cre).

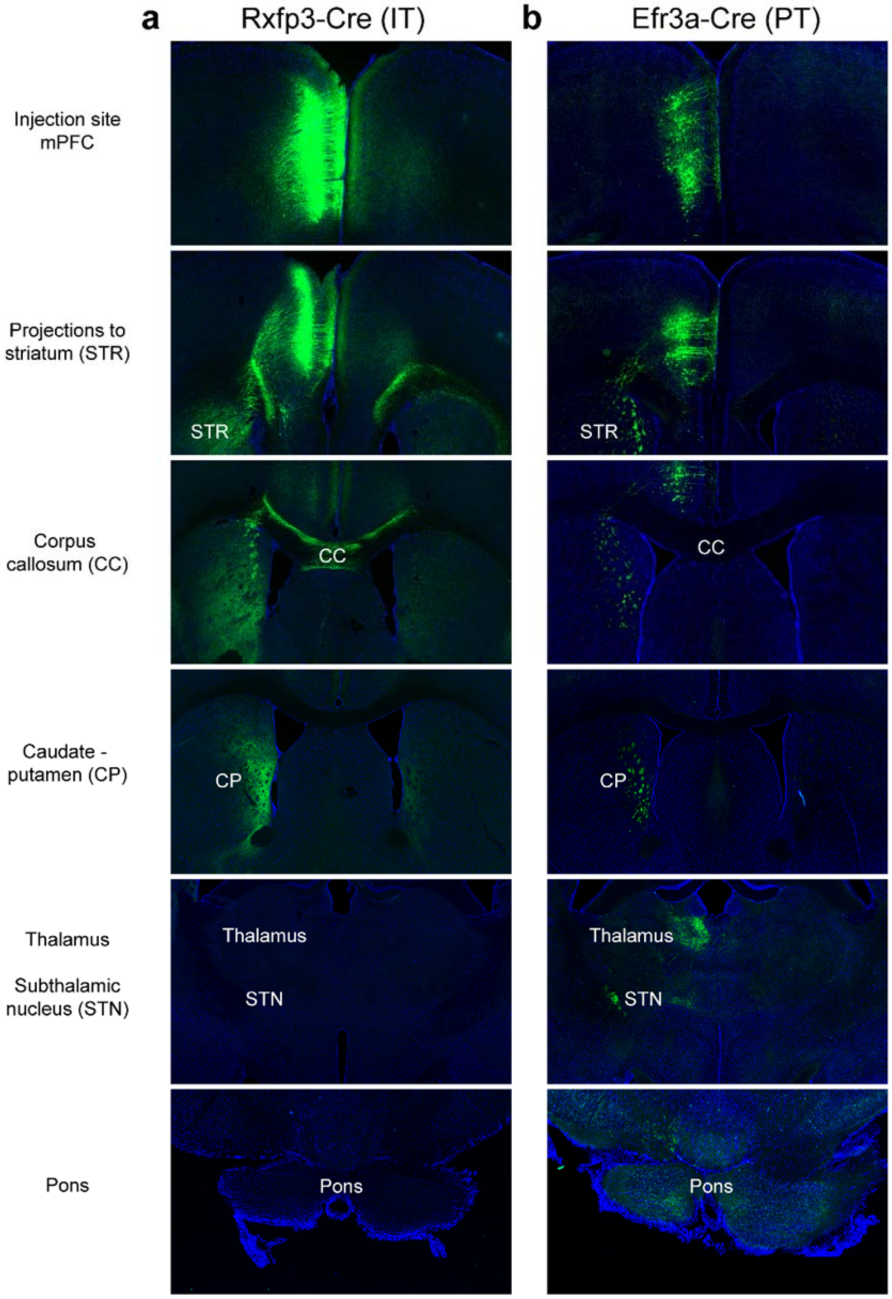


**Supplementary Fig. S2 | Histological verification of IT and PT neurons.** Shown are sample coronal brain sections showing fluorescence signals in Rxfp3-Cre and Efr3a-Cre mice. We injected AAV2/Ef1a-DIO-eYFP (in Rxfp3-Cre mouse) or AAV2-CAG-FLEX-tdTomato (in Efr3a-Cre mouse) unilaterally into the mPFC and performed histology four weeks later to examine the distribution of the fluorescence signals. As expected, fluorescence expression was detected in the deep layers of the mPFC in both mice. It was additionally detected in the contralateral cortices, as well as in the ipsilateral and contralateral striatums of Rxfp3-Cre mouse.

In Efr3a-Cre mouse, it was additionally detected in the ipsilateral striatum and other subcortical structures. **a,** In Rxfp3-Cre mouse, the injection labeled mPFC layer 5 neurons (AAV2-eYFP, green, top panel) that sent bilateral axonal projections to the striatum (STR, second to fourth panels) via the corpus callosum (CC, third panel), but not to the thalamus, subthalamic nucleus (STN) or pons (bottom two panels). This indicates selective labeling of IT neurons. **b,** In Efr3a-Cre mouse, the injection labeled mPFC layer 5 neurons that projected to the striatum, thalamus, and STN ipsilaterally (above pyramidal decussation), and to the pons bilaterally (below pyramidal decussation; AAV2-tdTomato, pseudo-colored green), indicating selective labeling of PT neurons.

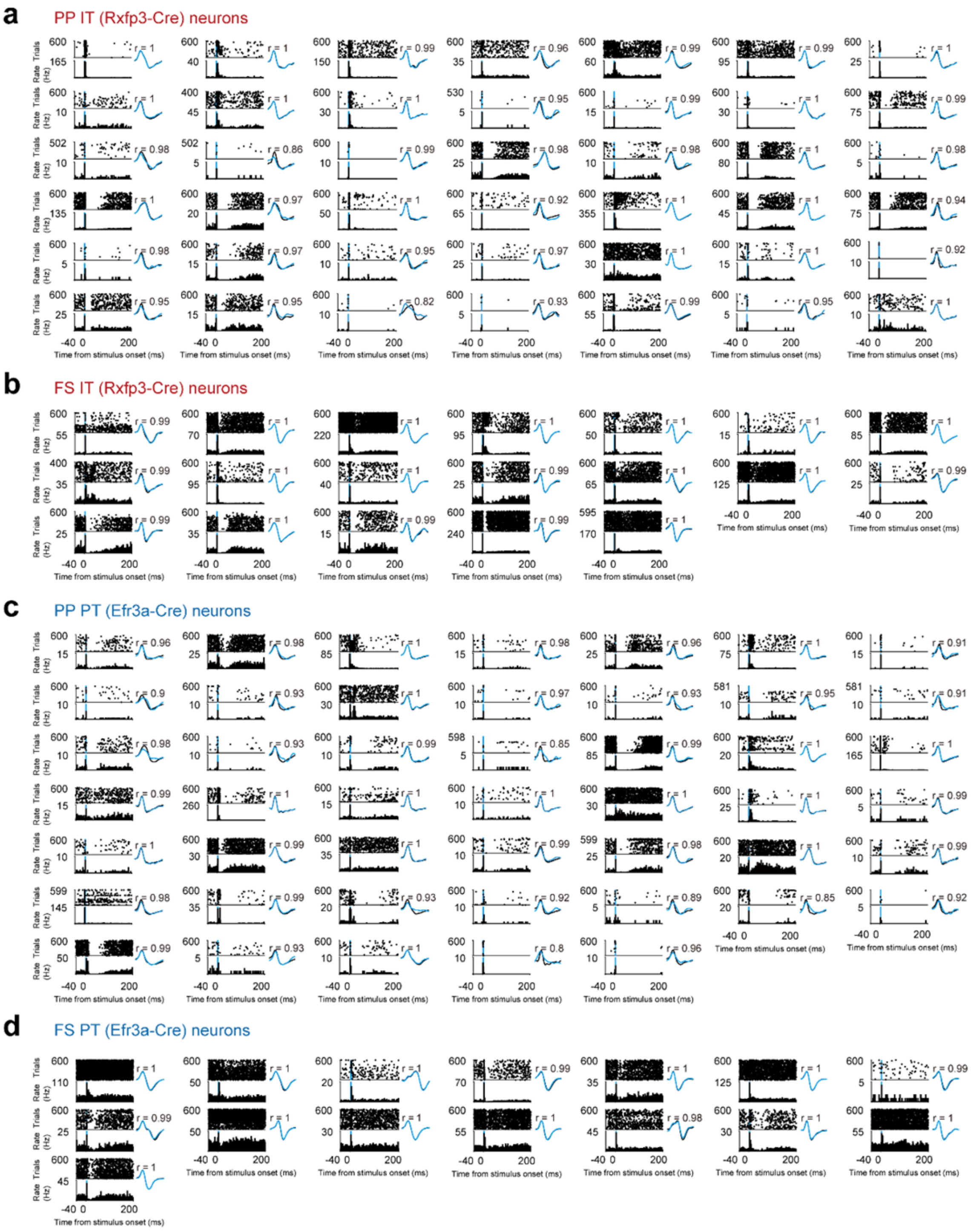


**Supplementary Fig. S3 | Optogenetic identification of IT and PT neurons.** Shown are the responses of optically tagged IT and PT neurons to laser stimulation (blue bars; 5 ms) recorded from Rxfp3-Cre (a, b) and Efr3a-Cre (c, d) mice, respectively. Optically tagged neurons classified neither as PP nor FS neurons (n = 5 from Rxfp3-Cre mice) are excluded. Top, spike raster plots; each line is one trial and each tick mark represents a spike. Bottom, peri-stimulus time histograms. Insets, averaged waveforms of spontaneous (black) and optically driven (blue) spikes (duration, 1 ms; calibration for spike amplitude varies across neurons; number, correlation coefficient between the two waveforms).

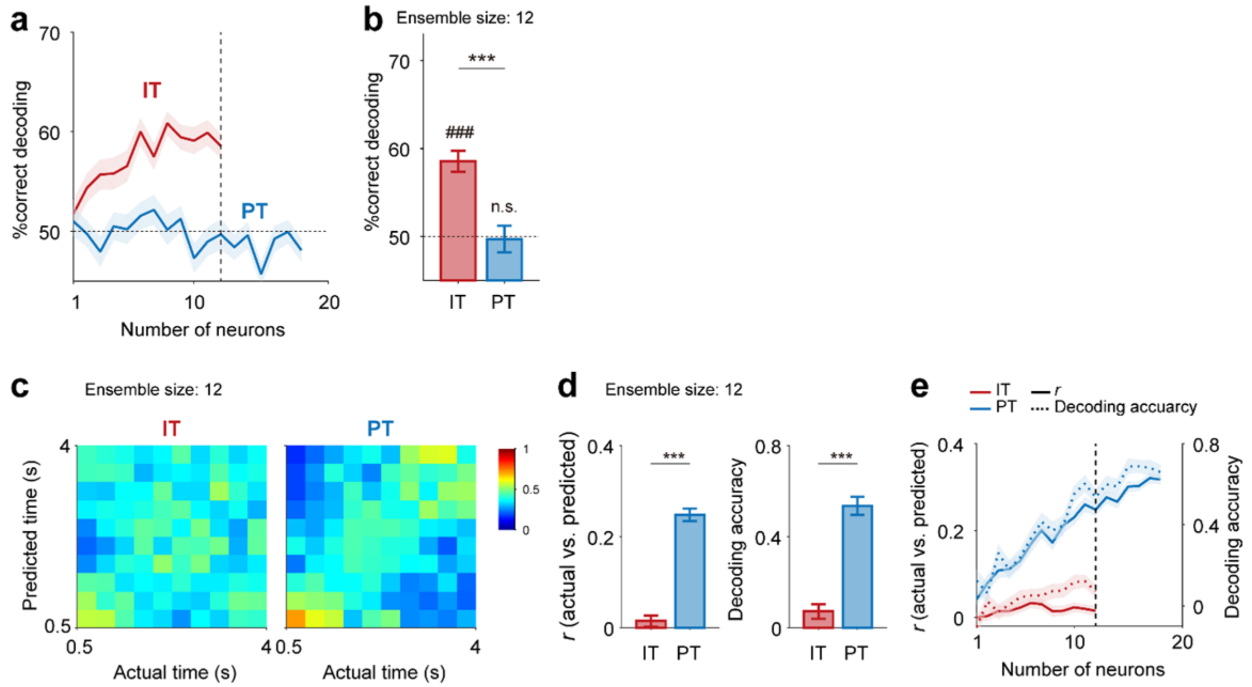


**Supplementary Fig. S4 | Results from the analysis of variable-delay trials.** We repeated the same analyses using the neural data (optically tagged PP neurons) collected during variable-delay trials. Neural data were analyzed up to 4 s using only those trials with durations ≥ 4 s. **a, b**, Working memory-related neural activity (ensemble size = 12 in b). Same format as in Fig. 3b, c. **c-e,** Timing-related neural activity (ensemble size = 12 in c, d). Same format as in Fig. 5a-c.

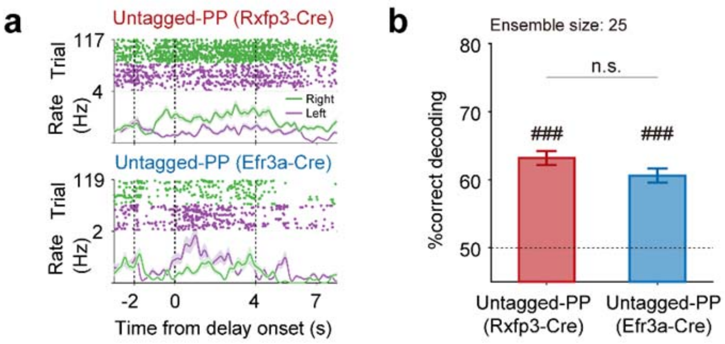


**Supplementary Fig. S5 | Comparison of working memory-related signals carried by untagged PP neurons between Rxfp3-Cre and Efr3a-Cre mice. a,** Sample responses of untagged PP neurons during the task (correct trials only). Top, spike raster plots; bottom, spike density functions ($\sigma$ = 100 ms). **b,** Decoding sample identity based on delay-period (4 s) ensemble activity of untagged PP neurons recorded from Rxfp3-Cre (red) or Efr3a-Cre (blue) mice (mean ± SEM across 100 iterations using 25 randomly selected neurons). ###$p < 0.001$ (above chance level, *t*-test); n.s., non-significant (between IT and PT, *t*-test).

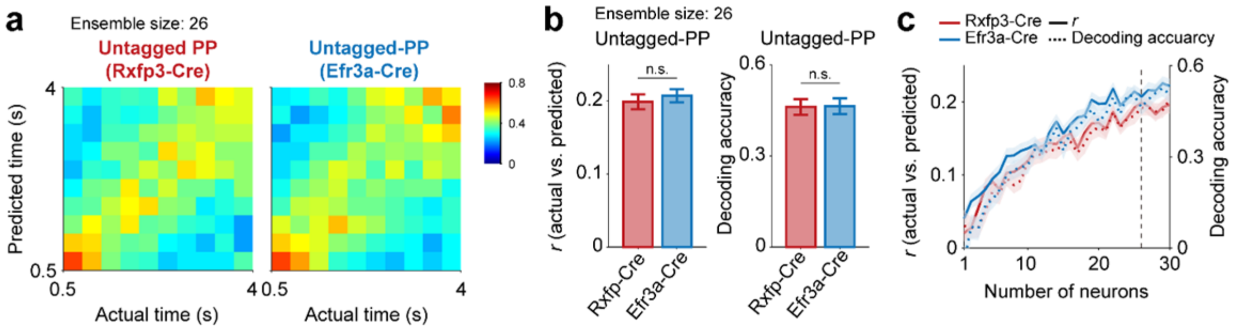


**Supplementary Fig. S6 | Comparison of temporal information carried by untagged PP neurons between Rxfp3-Cre and Efr3a-Cre mice. a,** Heat maps showing normalized decoding probabilities (actual versus predicted bins) using 26 randomly selected untagged PP neurons recorded from Rxfp3-Cre (left) or Efr3a-Cre (right) mice. **b,** Decoding performances (left, correlation between the actual and predicted bins; right, decoding accuracy; mean ± SEM across 100 decoding iterations of IT and PT neuronal ensembles (n = 26, vertical dashed line in c)). n.s., non-significant (*t*-test). **c,** Decoding performance as a function of ensemble size. Same format as in Fig. 5a-c.

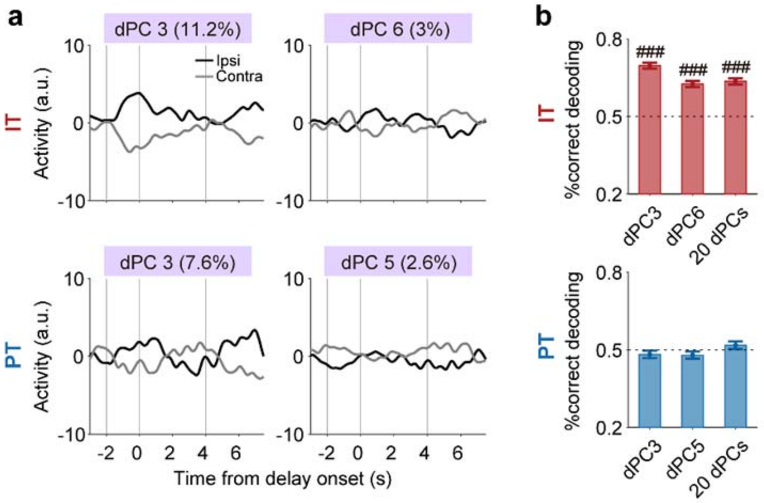


**Supplementary Fig. S7 | Sample-dependent dPCs of PT neurons do not convey significant working memory-related signals during the delay period. a,** Time courses of the top two sample-dependent dPCs. Black and gray denote ipsilateral and contralateral sample trials, respectively. **b,** Decoding performance (mean ± SEM across 100 decoding iterations) using only the delay-period component (4 s) of each sample-dependent dPC or 20 dPCs. ###$p$ < 0.001 (above chance level, $t$-test). These results indicate that sample dependence of these PT neuronal dPCs is because of sample-dependent neuronal activity before and/or after the delay period.
